# A framework for RNA quality correction in differential expression analysis

**DOI:** 10.1101/074245

**Authors:** Andrew E. Jaffe, Ran Tao, Alexis L. Norris, Marc Kealhofer, Abhinav Nellore, Yankai Jia, Thomas M. Hyde, Joel E. Kleinman, Richard E. Straub, Jeffrey T. Leek, Daniel R. Weinberger

## Abstract

RNA sequencing (RNA-seq) is a powerful approach for measuring gene expression levels in cells and tissues, but it relies on high-quality RNA. We demonstrate here that statistical adjustment employing existing quality measures largely fails to remove the effects of RNA degradation when RNA quality associates with the outcome of interest. Using RNA-seq data from a molecular degradation experiment of human brain tissue, we introduce the quality surrogate variable (qSVA) analysis framework for estimating and removing the confounding effect of RNA quality in differential expression analysis. We show this approach results in greatly improved replication rates (>3x) across two large independent postmortem human brain studies of schizophrenia. Finally, we explored public datasets to demonstrate potential RNA quality confounding when comparing expression levels of different brain regions and diagnostic groups beyond schizophrenia. Our approach can therefore improve the interpretation of differential expression analysis of transcriptomic data from the human brain.

## Introduction

Microarrays and RNA sequencing (RNA-seq) can measure gene expression levels across hundreds of samples in a single experiment. These experiments can be used to identify changes in gene expression associated with illness^1^, genetic variation^2^, drug treatment^3^, and development^4, 5^ and hundreds of these studies have been conducted in human tissue or cells, with a variety of outcomes^6^. As gene expression levels are measured with error, normalization procedures have been implemented for both microarray^7, 8^ and RNA sequencing^9, 10^ data to reduce technical variability, including controlling for variability associated with how and when the samples are run, so-called “batch” effects^11^. Recent research has further characterized this expression variability in RNA-seq data,^12–15^ including demonstrating variability associated with technical factors involved in the preparation, sequencing, and analysis of samples. Variability in gene expression is particularly influenced with RNA quality^16^ because accurately measuring gene expression levels strongly depends on the quality of the input RNA.

Many studies use postmortem human tissue as the source material, for example in the recent Genotype by Tissue Expression (GTEx) project, particularly for tissues and cells that are difficult to ascertain during life. Postmortem studies typically extract RNA from tissue that has been susceptible to a wide variety of antemortem and postmortem factors. Several approaches exist for quantifying the quality of the input RNA prior to sequencing library construction, including UV absorption ratios of 280nm to 260nm and RNA integrity numbers (RINs). RIN is a measurement largely derived from degradation profiles of ribosomal RNAs^17, 18^, and samples with low RINs can typically be excluded from data generation. Recommended RIN thresholds have been suggested as low as 5.0 for PCR^16^ and 7.0 for RNA-seq^18^. However, even high quality samples (RIN > 8) demonstrate evidence for degradation, as transcriptome-wide gene expression levels strongly associate with RIN even at higher RINs^15^. The recent introduction of ribosomal depletion approaches, like Ribozero library construction, has permitted the sequencing of lower quality samples compared to previous polyadenylation section-based approaches (polyA+) including samples with RINs less than 3.

The quality of RNA can also be estimated directly from RNA sequencing data, for example, by calculating the 5’ to 3’ read coverage bias (particularly in polyA+ data), transcript integrity numbers (TINs)^19^, various read mapping rates, including to autosomes, ribosomal RNAs, and mitochondrial RNAs (chrM), and gene/exon assignment rates^16^. While many of these approaches appear to capture the largest global effects on expression, for example through positively correlating factors of expression data with these quality measures, the presence and role of more subtle and gene/transcript specific biases in RNA quality on differential expression analysis of gene expression measurements is unclear.

Here we demonstrate that the existing RNA quality estimation approaches do not sufficiently correct for biased differential expression analysis when RNA quality differs or associates with the primary outcome of interest, particularly in the human brain. We have developed an experiment-based framework for RNA quality adjustment that resolves the deficiencies of earlier approaches. We first exposed human brain tissue to room temperature for varying amounts of time to generate a model landscape of molecular degradation in the human frontal cortex, using both polyA+ and RiboZero sequencing libraries. We then demonstrated that degradation time is more predictive of altered expression level measurements than traditional measures such as RIN. Using a large brain-derived polyA+ RNA-seq schizophrenia-control dataset (N=351), we demonstrate that the effect of degradation of each gene strongly and directionally predicts differential expression by case status. We further show that many existing approaches either fail or only partially reduce this degradation bias. We utilize the degradation RNA-seq data to identify transcriptomic regions more susceptible to degradation, and use the expression levels of these regions to correct for degradation bias. This approach, called quality surrogate variable analysis (qSVA), successfully removes degradation bias, and as evidence of its validity, improves replication of differential expression signals in independent RiboZero-based datasets. We also re-analyze two large human postmortem brain disease datasets that assayed differential expression with microarrays, as well as across tissues in the GTEX database, and find strong evidence of RNA quality confounding in each case that is corrected by qSVA. We therefore propose methods for assessing confounding by RNA degradation and a statistical framework - qSVA – for removing this degradation bias when present. These approaches can improve the interpretation of transcriptomic data from the human brain with potential applicability to other tissues.

## Results

### Different mRNAs degrade at different rates in human brain tissue

We first sought to characterize the transcriptional landscape of degradation in the human brain, given that most studies involving postmortem human tissue were obtained many hours after death and have a wide variety of RNA integrity numbers (RINs). We therefore left dorsolateral prefrontal cortex (DLPFC) tissue from five brains at room temperature (off of ice) for 0, 15, 30, and 60 minutes, extracted RNA, measured RINs, and then constructed and sequenced both polyA+ and RiboZero libraries (see Methods, Table S1). All five samples dropped over four units of RIN over just an hour off ice. After aligning and quantifying the RNA-seq data (see Methods), many technical covariates were strongly associated with degradation time in both library types, including RIN, read mapping rate, gene assignment rate, proportion of reads mapping to the mitochondrial chromosome (chrM), and the proportion of reads containing splice junctions (Table S2). Principal component analysis (PCA) suggested that degradation time was the strongest explanatory variable (PC1) in the expression levels across each library type, explaining 47.5% and 39.0% of normalized gene counts in polyA+ and RiboZero libraries respectively (Figure S1). Furthermore, degradation time was more predictive than RIN in each library type, as multivariate modeling retained the effect of degradation time but not RIN on PC1 (p=2.15×10^−4^ for polyA+ and p=8.52×10^−5^ for RiboZero). Linear modeling further demonstrated that many genes were highly susceptible to the effects of RNA degradation, including 12,324 genes at FDR < 5% significance in the polyA+ dataset (N=2,303 at p_bonf_ < 5%) and 10,981 genes in the RiboZero dataset (N=2017 at p_bonf_ < 5%, Table S3).

Gene set analysis on the degree of degradation suggested dysregulation of a wide variety of cellular processes (Table S4). Among the 211 gene sets with FDR significance (< 5%) among the degradation test statistics from both library types, all but two in polyA+ (related to ribosomal subunits) were susceptible to degradation (rather than protective). Furthermore, as RNAs from different cell types may degrade at different rates, we compared the degree of degradation to the degree of cell type-specificity from single cell sequencing^20^. Those genes most differentially expressed across cell types in the human brain were significantly more likely to be associated with degradation (Figure S2). For example, the most cell type-specific genes (F > 10, N=2640) had degradation t-statistics on average 3.33 units lower in RiboZero (p < 10^−100^) and 1.64 units lower in polyA+ (p=1.22×10^−88^) data compared to those genes not associated with cell type (F<1). Similarly, comparing the most cell type-specific genes for each cell type more directly suggested differential susceptibility to degradation (see Methods), with a larger effect in RiboZero data (Table S5). For example, in the RiboZero data, those genes marking astrocytes (N=224 genes, ΔT= −42.44, p=1.12×10^−41^) and endothelial cells (N=190, ΔT= −3.70, p=14.77×10^−41^) were most susceptible to degradation followed closely by neurons (N=244, ΔT= −2.83, p=3.26×10^−31^). Genes specific for other cell types appeared less susceptible to degradation, including oligodendrocytes (N=222, ΔT= −2.13., p=6.16×10^−17^) and microglia (N=186, ΔT= −1.77, p=1.77×10^−9^), whereas oligodendrocyte progenitor cells appeared unaffected by degradation (N=126, ΔT= −0.65, p=5.41×10^−2^). In PolyA+ data, genes marking neurons were most susceptible to degradation (N=241, ΔT= −1.75 p=4.41×10^−10^), followed by astrocytes (N=222, ΔT= −1.01, p=5.67×10^−4^). In contrast to RiboZero data, genes marking endothelial cells appeared resistant to degradation in the polyA+ data (N=196, ΔT= 0.82, p=8.57×10^−3^). These enrichment analyses indeed suggest that RNAs from different cell types may be differentially susceptible to degradation, which is captured uniquely by different RNA-seq library preparation methods.

We next examined whether these degradation effects were brain- and degradation method specific. Our degradation data was compared to polyA+ RNA-seq data from a similarly designed blood degradation dataset, where blood samples from 4 individuals were left at room temperature at 0, 12, 24, 48, and 84 hours, and then sequenced using the polyA+ protocol^21^. The rate of degradation, as measured by RIN, was more rapid in our brain samples, as blood samples still had RINs > 7.7 after 12 hours at room temperature, compared to our brain samples having RINs under 6.6 after just one hour. We also find only weak global correlation between the effect of degradation on blood and frontal cortex gene expression (Figure S3A), and much smaller degradation rates of individual genes in blood compared to brain (median: 33.6% versus 0.44%, 90^th^ percentile: 213.8% versus 1.4% change per hour). Conversely, we found that our tissue degradation experiment showed relatively similar effects to that of RNA degradation based on treating RNA with RNase A. Using the 12 samples in the ABRF study that used Illumina sequencing of human brain reference RNA with the Ribozero protocol^22^, we compared the 3 samples treated with RNase A to the 9 untreated samples. We found 13,553 genes significantly associated with RNase treatment (at FDR < 5%), of which 7,700 (65.7%) were also significantly differentially expressed in our tissue degradation experiment, a 6.28 fold enrichment. Globally, the resulting degradation effect T-statistics were highly correlated (Figure S3B), suggesting that RNase A-like activity is a major factor contributing to degradation in postmortem human brain tissue following death.

### Strong bias in differential expression analysis in confounded designs

We thus reasoned that many results in differential expression analyses of postmortem brain datasets may be susceptible to RNA degradation confounding. For example, many studies comparing different diagnostic groups typically have significant group differences in measures of RNA quality (e.g. RINs). We therefore utilized two large RNA-seq datasets from the prefrontal cortex comparing patients with schizophrenia to adult controls: Lieber Institute for Brain Development (LIBD, “discovery” data, polyA+ protocol, N=351) and CommonMind Consortium (CMC, “replication” data, RiboZero protocol, N=331) – see companion Jaffe et al 2016 for details. We first create a new diagnostic plot to compare differential expression statistics for diagnosis in these datasets to the degradation statistics from our RNA degradation experiment (fold change in expression per minute, or its corresponding t-statistic). This approach, which we call the “differential expression quality” (DEqual) plot, can quantify the degree of potential bias by RNA degradation across the entire transcriptome in a given dataset. Using these DEqual plots, we observed strong positive correlation between univariate differential expression statistics for diagnosis and ex vivo degradation in the LIBD dataset (Figure 1A). This plots shows that genes positively associated with degradation time (e.g. are relatively protected during degradation) are almost always more highly expressed in the outcome group compared to the control group, and genes negatively associated with degradation time were almost uniformly more highly expressed in the outcome group. Thus the directionality of change associated with an outcome at a particular gene can be predicted almost entirely by its relationship with degradation and the difference in RNA quality between outcome groups. Similar degradation associations are present using log_2_ fold changes rather than t-statistics (Figure S4A). In both the LIBD and CMC datasets, the patient group had lower RINs than controls (LIBD: p=4.48×10^−5^, CMC: p=8.63×10^−7^), and among the 24,122 genes with RPKM > 0.1, we found that 11,408 (47.3%) were significantly differentially expressed at FDR < 5%, further suggesting confounding by RNA degradation. The degree of correlation in these plots can therefore be a useful quality control metric for potential degradation bias in differential expression analysis.

**Figure 1:**
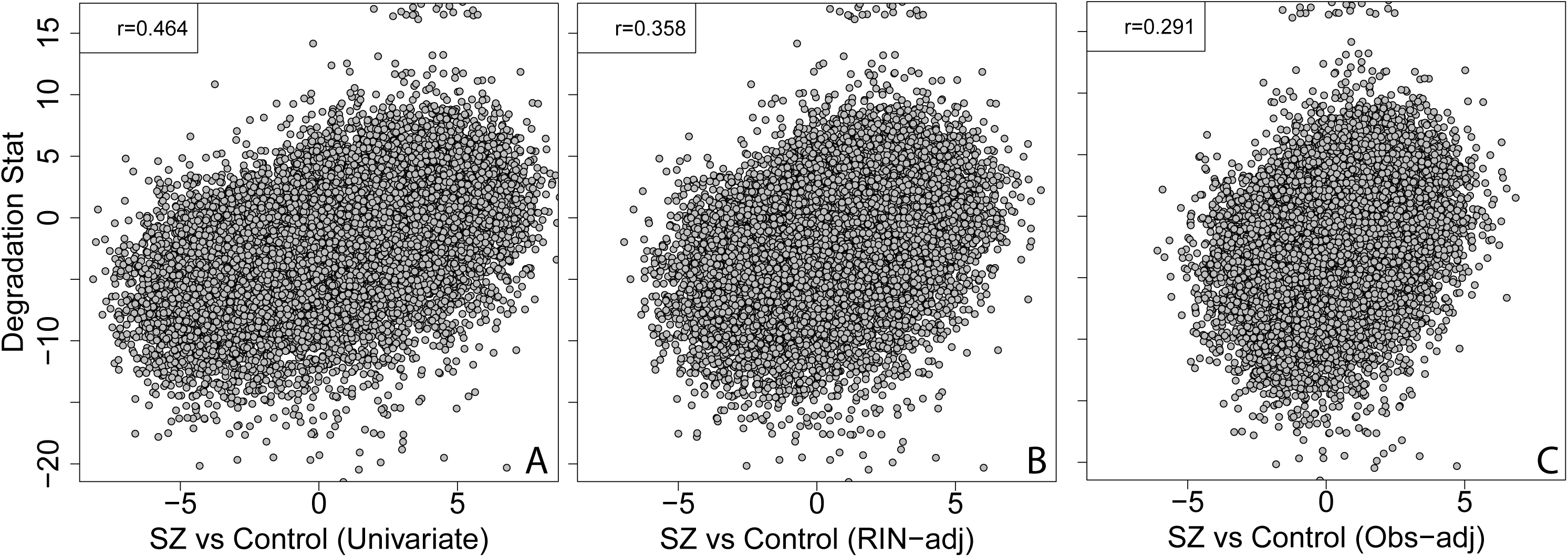
Differential expression quality (DEqual) plots for schizophrenia-control expression differences. Each DEqual plot compares the effect of RNA degradation from an independent degradation experiment on the y-axis from to the effect of the outcome of interest, here schizophrenia (SCZD) compared to controls. Each point is a gene, and effects here are shown as T-statistics for each effect. (A) DEqual plot for univariate case-control analysis shows strong correlation between degradation and diagnosis effects, (B) DEqual plot for RIN-adjusted case-control differences largely fails to remove degradation bias, and (C) DEqual plot when adjusting for observed clinical and technical covariates, including age, sex, ethnicity, chrM mapping rate, gene assignment rate, and RIN, also fail to remove degradation bias.

### Adjusting for RIN fails to remove degradation bias

Given the DEqual plots from the univariate analysis, the significant difference in RIN between the schizophrenia and control groups, and the large number of differentially expressed genes, we expected that adjusting the differential expression model for RINs would reduce the degree of degradation bias. However, RIN adjustment only partially reduced the correlation between diagnosis and degradation statistics (Figure 1B), from ρ = 0.464 to ρ=0.358, and only reduced the number of FDR-significant differentially expressed genes from 11,408 to 6,622. The degree of RNA degradation bias was practically identical when further modeling RIN non-linearly, e.g. further adjusting for RIN and RIN^2^ (Figure S5). We further adjusted the differential expression analysis for other observed variables, including clinical and technical covariates (“observed” model: age, sex, ethnicity, chrM map rate, gene assignment rate, and RIN), which again only partially reduced the correlation between diagnosis and degradation statistics (Figure 1C, to ρ=0.291) and the number of genes (N=2,215) significantly differentially expressed.

We also used the blood degradation dataset to demonstrate that RIN adjustment fails to account for the true degradation effect. Here, we modeled differences in expression between individuals 1 and 2 after inducing confounding by degradation time (the true degradation effect) by removing one time point per individual (see Methods). As expected, the DEqual plot in univariate analysis shows strong correlation between the individual effect and the degradation effect (Figure 2A, ρ=0.495). We can again show using the DEqual plot that the RIN adjustment does not remove the strong degradation bias (Figure 2B, ρ = 0.307). Here, in this dataset, unlike the schizophrenia case-control datasets described above, we have measured the true degradation variable – time at room temperature – and show that adjusting for this measure completely removes the RNA degradation bias (Figure 2C, ρ=-0.09). These results suggest that RIN or other observed quality variables maybe be a poor surrogate for total RNA quality and that adjusting for RIN in differential expression analysis is insufficient to remove potential RNA degradation bias.

**Figure 2:**
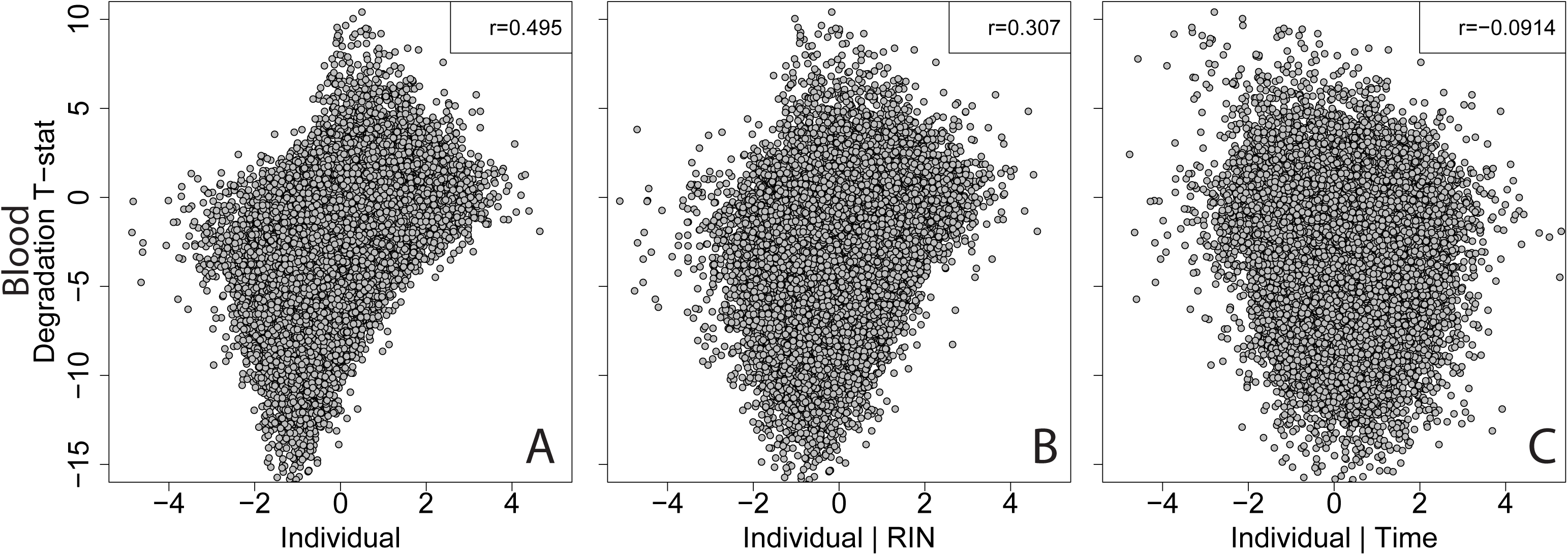
RIN adjustment largely fails to remove RNA quality bias. DEqual plots in blood degradation dataset after inducing confounding suggest RIN-adjustment fails to remove degradation bias. (A) DEqual plot for the effect of individual on expression after inducing confounding by degradation time shows strong bias. (B) DEqual plot for the individual effect after adjusting for RIN largely fails to remove degradation bias. (C) Adjusting for the true degradation variable – time – successfully removes degradation bias in the DEqual plot.

### Quality Surrogate Variable Analysis (qSVA) corrects RNA degradation bias

We hypothesized that we could leverage the degradation DLPFC RNA-seq data to better estimate factors related to RNA quality in postmortem expression datasets. This approach relied on estimating the genomic regions most susceptible to RNA degradation and using these regions as “negative control” features akin to approaches such as Remove Unwanted Variation (RUV)^10^ or Surrogate Variable Analysis (SVA)^23^. Specifically, we used the tissue degradation RNA-seq data to identify the expressed regions (ERs)^24^ most susceptible to RNA degradation (see Methods), resulting in 1000 “degradation regions” (chr:start-end) most susceptible to degradation in polyA+ libraries and 515 degradation regions most susceptible to degradation in RiboZero libraries. The regions across these two library types were completely non-overlapping, suggesting that different library protocols are susceptible to different RNA degradation profiles. The correction algorithm consists of quantifying coverage of these regions in datasets to be corrected for RNA quality to form the region by sample “degradation matrix”. Then factor analysis on the log-transformed degradation matrix generates “quality surrogate variables” (qSVs) that can be included as adjustment variables in differential expression analysis. The number of qSVs to include in analysis can be estimated using permutation-based approaches, for example using the Buja-Eyuboglu algorithm^25^ or using other visual approaches for selecting the appropriate number of factors to include in analysis. The qSVA approach is now available in the sva Bioconductor package^26^ and example code to run the correction procedure is described in the Methods section.

We applied the qSVA algorithm to the LIBD polyA+ RNA-seq data with the “observed” model (consisting of observed clinical and technical confounders) described above. Here, the DEqual plot showed completely attenuated degradation bias (Figure 3, ρ=-0.11 using t-statistics and ρ=-0.037 using log_2_ fold changes). This change in directionality in the DEqual plots was very similar comparing the RIN-versus time-adjusted modeling in the blood dataset (see Figure 2), where the true confounding variable (time) was known. Here there were also only 183 genes differentially expressed at FDR < 5% further suggesting a reduction of RNA degradation bias in differential expression analysis of schizophrenia patients versus controls. The qSVs themselves were strongly associated with observed variables including chrM alignment rate, RIN, total gene assignment rate, overall alignment rate, age, and postmortem interval (see Figure S6). Similarly, in the CMC dataset, the qSVs, calculated using the RiboZero-based degradation regions, were strongly associated with RIN, total gene assignment rate, institute where the sample was collected, and sequencing and flow cell batches (Figure S7). These results suggest that enriching for degradation signal via the independent tissue degradation experiment can identify more robust measures of RNA quality directly from RNA-seq experiments than relying on single observable measures.

**Figure 3:**
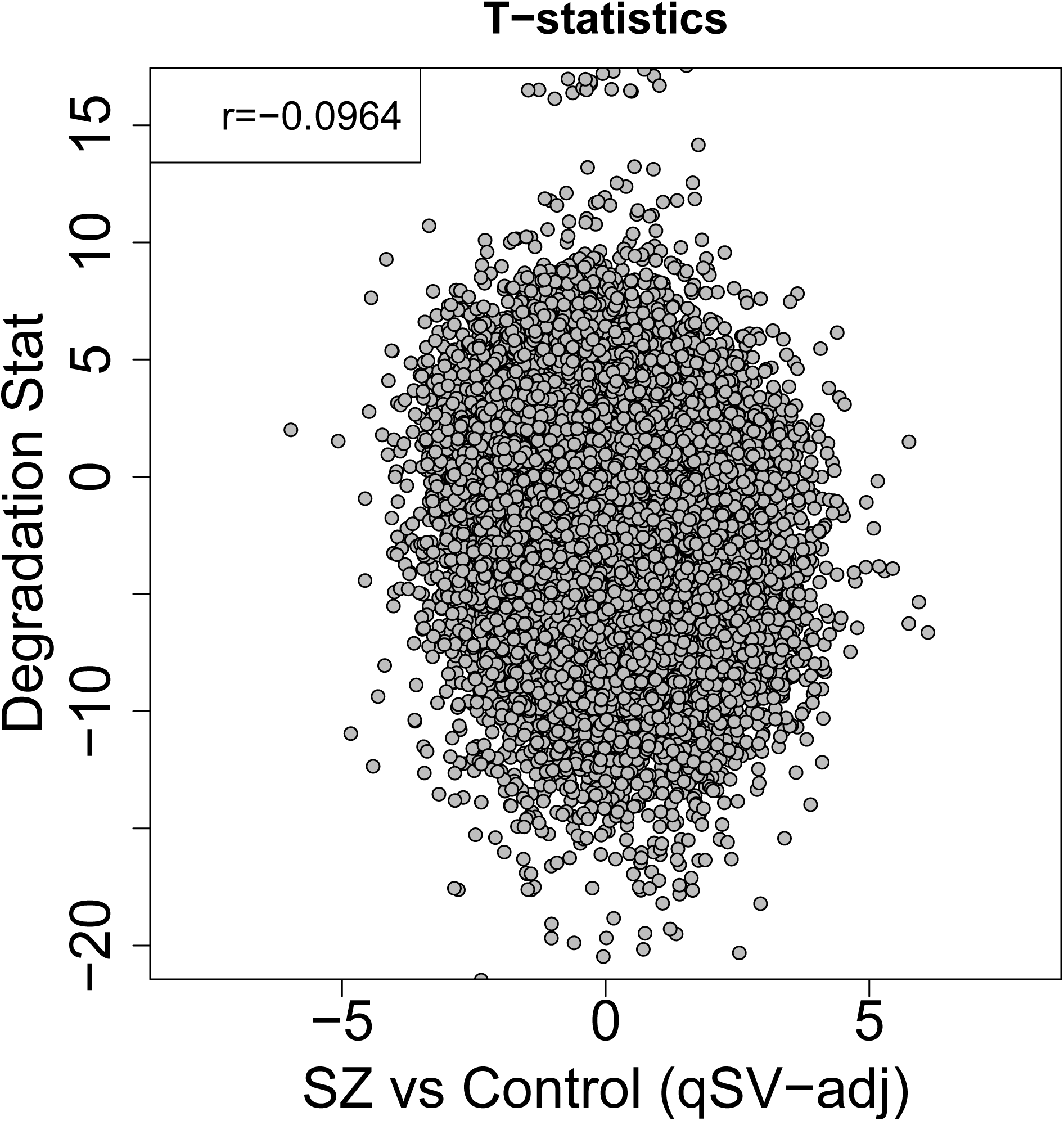
DEqual plot demonstrating that the quality surrogate variable (qSVA) framework successfully removes positive correlation between degradation and SCZD effects.

**Figure 4:**
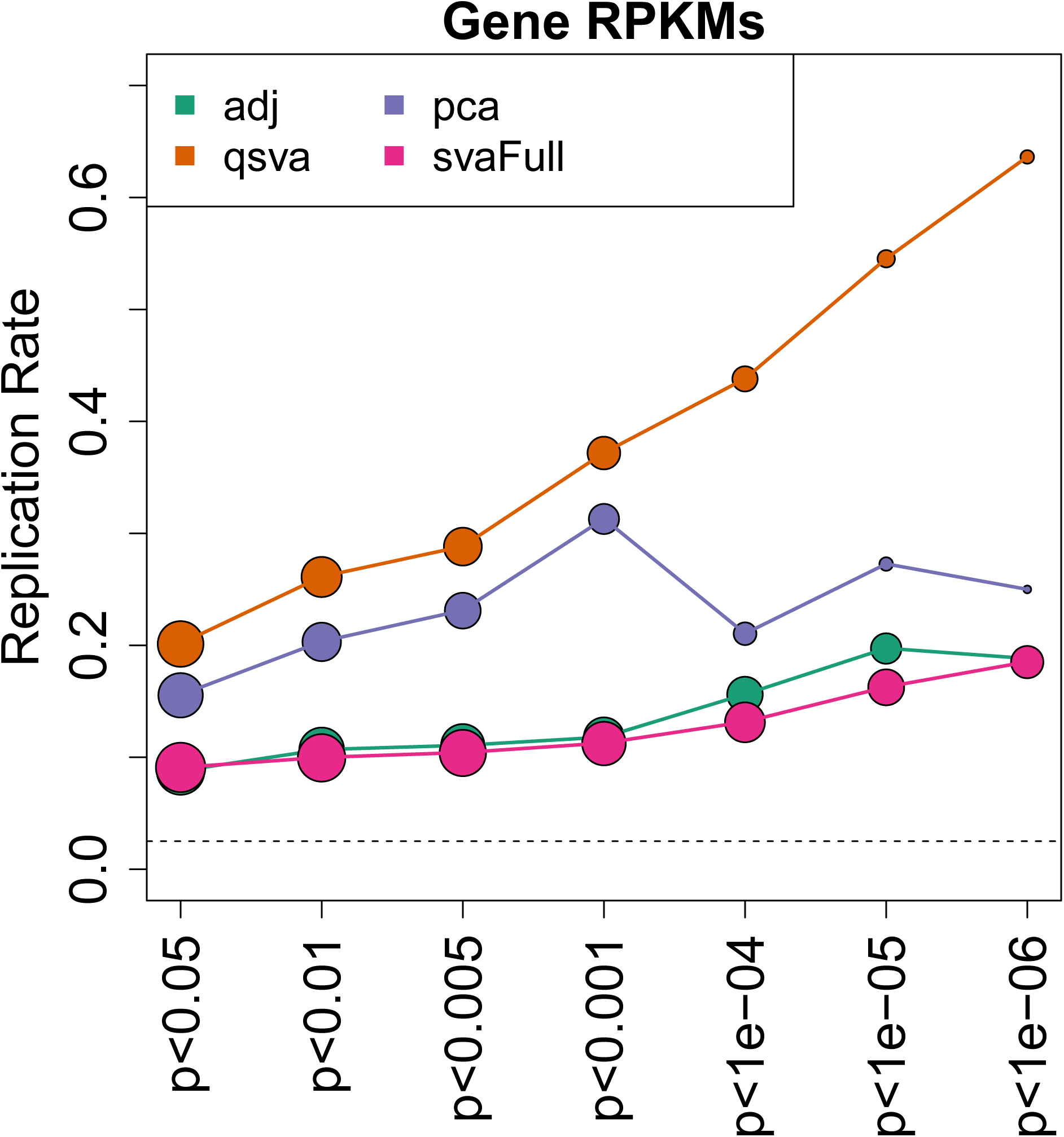
qSVA improves replication across independent datasets. We modeled SCZD-control expression differences using four statistical models in the LIBD (discovery) and CMC (replication) datasets. For a given significance threshold in the discovery dataset, we computed the replication rate (same fold change direction for case status and p < 0.05) in the replication dataset. The qSVA approach had the highest replication rate, while the covariate-adjusted and SVA approaches had the lowest replication rates.

### Improved replication for schizophrenia differential expression using qSVA

While the qSVA approach appears to remove RNA degradation bias in brain differential expression analysis via the DEqual plot, we further observed that adjusting for transcriptome-wide principal components (PCs) also removes the degradation effects in the DEqual plot (Figure S8, ρ =-0.02). This suggests that factor-based approaches – including qSVA but also more generally PCA - can identify and subsequently remove latent measures of RNA quality. However, “supervised” factor based approaches, like surrogate variable analysis (SVA) that rely on residualizing around a provided statistical model, largely preserved RNA degradation bias as shown in the DEqual plots (Figure S9).

We used replication signal across independent datasets – LIBD and CMC – to more fully contrast the value of the different degradation adjustment approaches. For a given adjustment approach, we calculated replication rates of differentially expressed genes discovered in the LIBD dataset at different significance thresholds in the CMC dataset. We found the lowest replication rates (<20%) regardless of significance threshold when adjusting only for observed clinical and technical variables including RIN, as well as SVA residualizing on only diagnosis.

While we had high replication rates among marginally significant genes (p<0.001) using SVA residualizing on the observed variables described above, we found strong inflation of the test statistics among both the LIBD (9,033 genes at FDR < 5%) and CMC (6,924 genes at FDR < 5%) datasets. Among those genes significantly differentially expressed (p<10^−4^), we found the highest replication rates using qSVA, as well as relatively linear improvements in the replication rate as the discovery p-values threshold dropped. Importantly, the qSVs calculated in the LIBD and CMC datasets were based on different degradation regions, as the CMC data was RiboZero and the LIBD data was polyA+. These results therefore show that qSVA leads to improved replication in postmortem brain transcriptomic studies.

### Applicability of qSVA to other tissues and brain regions

We next calculated the relevance of the degradation regions derived from human brain tissue to the blood and ABRF RNase A brain RNA degradation datasets. After performing the qSVA algorithm using the relevant library type degradation regions (polyA+ for blood, RiboZero for commercial brain RNA), we found high predictive accuracy in both datasets. For example, in the blood dataset, the first qSV was strongly associated with degradation time (p=7.93×10^−9^, Figure S10A), as was the first qSV in the brain commercial RNA dataset (p=3.57×10^−7^, Figure S10B). In the confounded individual example from the blood dataset, we can successfully remove degradation bias, as illustrated in the DEqual plot, by adjusting for the first qSV based on the brain degradation data (Figure S11A). More importantly, the qSV adjustment results in more statistically unbiased effect estimates (i.e. log_2_ fold changes for the “individual” effect) compared to the statistical model adjusting for degradation time, which is the true confounding variable (Figure S11B). Conversely, the statistical bias in differential expression signal from the RIN-adjusted model for the “individual” effect relative to the degradation time-adjusted model is much larger (Figure S11C). These results suggest that perhaps general signatures of degradation that translate to other tissues can be estimated using human brain tissue RNA-seq degredation data, and potentially be used to correct RNA degradation bias in other tissues.

We also defined the top 1000 degradation susceptible regions based on an analogous procedure in the blood RNA-seq data. There were only 4 regions in common between blood and DLPFC degradation-susceptible regions (within genes: *PNKD*, *MBOAT7*, *ENG*, and *SULF2*).

We performed the qSVA algorithm using these blood degradation-susceptible regions in the LIBD SZ-control dataset, and evaluated the performance using DEqual plots and calculating the number of genes significantly differently expressed. Here while the log_2_ fold changes when adjusting using blood versus brain degradation regions were correlated (Figure 12A, ρ=0.6), there was stronger negative correlation between degradation susceptibility in brain and blood-adjusted case control differences (Figure S12A, ρ=-0.11). However, using the blood degradation qSVA yielded 1,057 genes significantly differentially expressed at FDR < 5%, approximately five times more than using the brain degradation-susceptible regions.

We next used GTEx RNA-seq expression data - N=9,502 across 49 detailed tissues - to characterize the generalizability of DLPFC-derived degradation regions to other brain regions and tissue types. We compared each of 48 detailed tissues in GTEx to BA9 frontal cortex (the 49^th^ detailed tissue) using differential expression analysis before and after qSVA correction. In the univariate modeling for each tissue versus DLPFC, we found strong association between resulting correlation in DEqual plots and the difference in chrM mapping rates (Figure 13A, ρ=0.736, p= 2.44×10^−9^). These quality associations were driven by the 12 other brain regions (ρ=0.88, p= 2.44×10^−9^) as the non-brain tissues showed no association (ρ=0.19, p=0.26). Here qSVA correction removed the overall quality effects across the sub-tissues, largely by removing the positive correlation in the brain samples (Figure 13B, ρ=0.0, p=0.97). These results suggest that using DLPFC-derived degradation regions for qSVA correction may work well in other brain regions, but may not be appropriate for RNA degradation correction in other tissues in the body (see Discussion).

### Degradation bias signal in published differential expression analyses

We lastly compared the presence of RNA quality bias in published differential expression analyses in human brain for different disorders. As there are currently few additional large RNA-seq studies of postmortem human brain tissue in disease states, we used previously published large microarray datasets on differential expression in autism spectrum disorder (ASD)^27^ and Alzheimer’s disease (AD)^28^ across multiple brain regions. In the ASD dataset, patients had significantly lower RINs than controls in the frontal (p=0.021) but not temporal (p=0.70) cortex, and in the AD dataset, patients were significantly lower than controls for the single RIN provided across the three brain regions (p=1.23×10^−10^). After using all the probes on each microarray platform as surrogates for RNA-seq reads and mapping them to the genome, we quantified their corresponding degradation susceptibility based on our RNA degradation data. We selected those probes that were most significantly associated with degradation akin to our procedure described above for RNA-seq data (see Methods). In the ASD dataset, we found that those probes most associated with degradation (N=1,129 at p_bonf_ < 1%) were almost uniformly more lowly expressed in patients compared to controls in the frontal cortex (Figure 5A, p=2.2×10^−49^). The directionality of enrichment was consistent between diagnosis and degradation associations, given that almost all degradation susceptible probes decreased in expression over time (98.5%) and RINs were lower in patients compared to controls. In the temporal cortex, where RINs did not differ between cases and controls, there was attenuated enrichment in the same negative direction (p=4.77×10^−6^). Following the qSVA procedure (PCA on the 1129 susceptible probes and the adjustment for the resulting qSVs), the association between degradation-susceptible probes and diagnosis was removed (p=0.496, Figure 5B).

**Figure 5:**
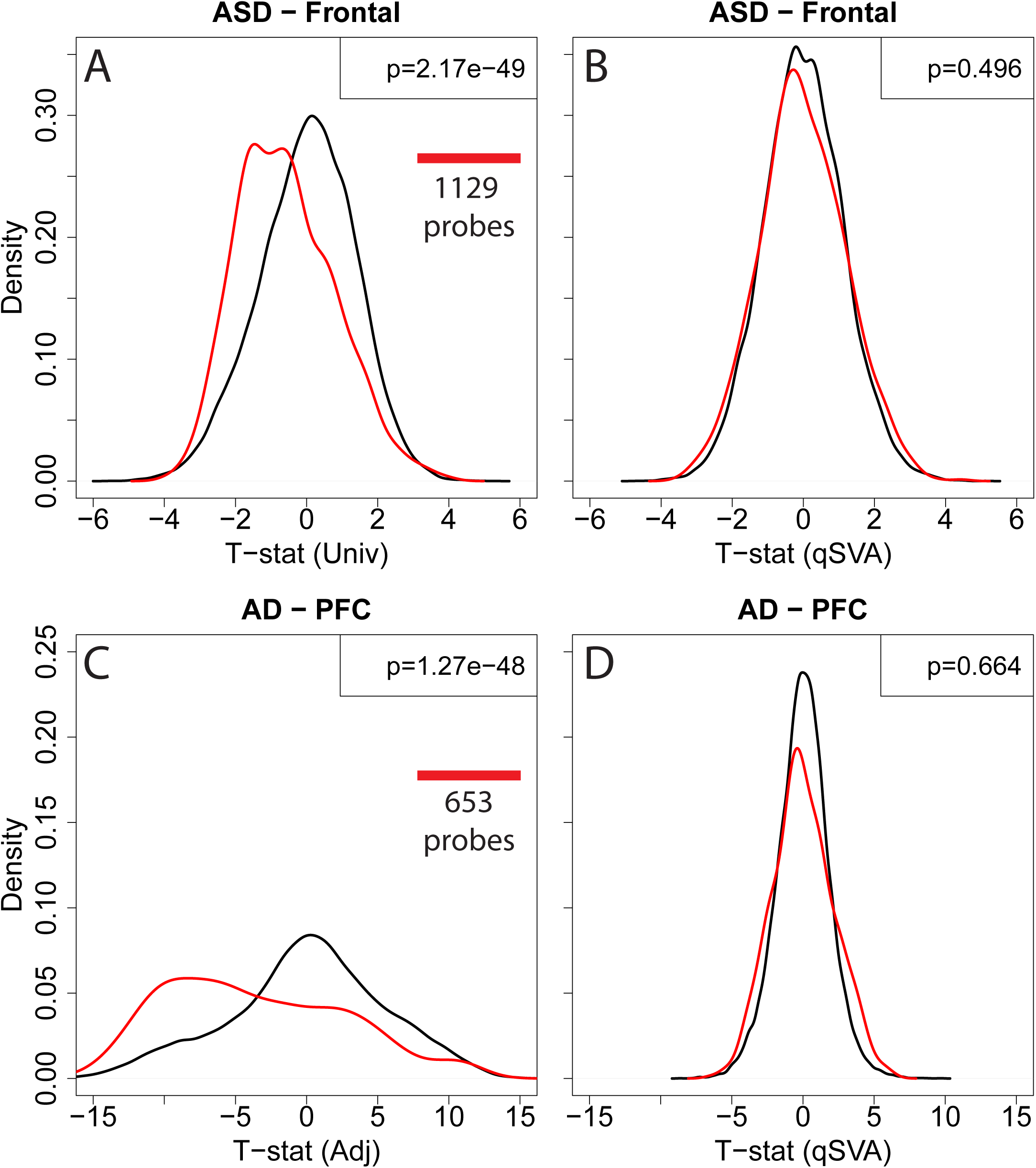
Differential expression signal for other diagnoses associated with RNA degradation. (A) Densities of the autism spectrum disorder (ASD) univariate effect in the frontal cortex, stratified by whether microarray probes were significantly associated with RNA degradation. (B) Stratified densities of qSVA-adjusted ASD effect suggest no residual association with RNA degradation. Analogous plots for Alzheimer’s disease (AD) for (C) covariate (age, sex, and batch) adjusted and (D) qSV-adjusted AD effect stratified by whether each probe significantly associated with degradation.

We found the same enrichment among differentially expressed probes for AD across all 3 brain regions and the 653 degradation-susceptible probes on this microarray, including in the prefrontal cortex (p=1.27×10^−48^, Figure 5C), cerebellum (p=1.82×10^−33^) and visual cortex (p=2.35×10^−35^). Adjusting for the resulting qSVs again removed the association between diagnosis and degradation susceptibility in the prefrontal cortex (p=0.66, Figure 5D) and cerebellum (p=0.49), and greatly reduced the association in the visual cortex (p=6.11×10^−4^). The qSVA correction also greatly reduced the magnitude of the differential expression test statistics across the entire platform (Figure 7C versus Figure 7D). These results further underscore the risk of potentially spurious findings based on uncorrected RNA quality confounding.

## Discussion

In this report, we characterized aspects of the landscape of RNA degradation across the human DLPFC transcriptome and identified largely brain-specific degradation signals. The cell types represented in homogenate brain tissue further showed differential susceptibility to RNA degradation. We used this DLPFC degradation data to identify the most degradation-susceptible regions in polyA+ and RiboZero RNA sequencing libraries, and developed an approached called qSVA to estimate and remove RNA degradation bias in differential expression analyses. We showed that the qSVA approach results in better replication across independent studies and in various public tissue datasets than existing popular statistical models that model observed measures of RNA quality like RIN, chrM mapping rate, and gene assignment rate. Our qSVA approach has a potential advantage over general PCA or RUV adjustments-less risk of removing true signals along with the noise. Re-analysis of previously published microarray datasets for Alzheimer’s disease (AD) and autism spectrum disorder (ASD) further suggested that probes differentially expressed for diagnosis were highly associated in a predictable directionality with RNA degradation susceptibility in both datasets.

We demonstrate that adjusting for measures of RIN largely fails to remove RNA degradation bias, and formally show that RIN correction is more statistically biased at estimating fold changes than qSVA when the true degradation effect is known. We hypothesize that RIN correction fails to remove these degradation bias effects for several reasons. The estimation of RIN itself is heavily driven by the intactness of ribosomal RNAs^17^, which appears only weakly associated with the underlying quality of total or polyadenylated RNAs across different subjects or tissues. Similarly, total RNA quality may be more complex than a single number per sample, as the resulting qSVs in both the LIBD and CMC datasets associate with a variety of technical factors (Figures S6 and S7) that may each influence RNA quality in subtle ways. Therefore, while the RIN value may be a rough guide in determining whether or not to sequence a particular sample, we would argue based on the data here, that it is not a particularly accurate or useful gauge of RNA quality.

The applicability of DLPFC-derived degradation-susceptible regions to other brain regions or tissues in the body is an important consideration in differential expression analysis. One practical recommendation for other brain regions would be use the degradation data from DLPFC to create DEqual plots, quantify the potential RNA degradation bias from its correlation and then evaluate how the DEqual plot changes when performing qSVA using the DLPFC degradation regions. If this qSVA correction fails to remove strong correlation between differential expression effects of degradation and outcome, researchers can also generate their own reference degradation datasets and apply the qSVA algorithm.

These results highlight the need to more carefully consider RNA quality as a potential confound in differential expression analysis, particularly when RNA quality associates with the outcome of interest. We argue that differences in latent RNA quality and the underlying cellular composition of homogenate tissue sources^29–31^ are two of the strongest confounding factors in postmortem human studies. The qSVA approach here that uses quality-associated features is analogous to our previously proposed approach that uses cell type-associated features to untangle the confounding effects of cellular composition (sparse PCA)^32^. The current study does suggest a potential interaction between RNA quality and cellular composition (Figure S2, Table S5) which may be more difficult to statistically isolate the two strong confounding effects. Nevertheless, our approach for directly measuring and statistically modeling RNA quality can improve the interpretation of differential expression analysis of transcriptomic data from the human brain.

## Methods

### Postmortem brain samples

Post-mortem human brain tissue was obtained by autopsy primarily from the Offices of the Chief Medical Examiner of the State of Maryland all with informed consent from the legal next of kin (Protocol # 12-24 approved by the Institutional Review Board of the Department of Health and Mental Hygiene of the State of Maryland). Clinical characterization, diagnoses, and macro- and microscopic neuropathological examinations were performed on all samples using a standardized paradigm, and subjects with evidence of macro- or microscopic neuropathology were excluded. Details of tissue acquisition, handling, processing, dissection, clinical characterization, diagnoses, neuropathological examinations, RNA extraction and quality control measures were described previously in Lipska, et al.^33^.

### Tissue degradation experiment

DLPFC gray matter from 5 donors was dissected, pulverized and mixed on dry ice, and then start the experiment: aliquot ~100mg pulverized tissue for 4 times for each subject on dry ice and then put tissue aliquots on room temperature except 1 aliquot of each subject, which is the kept on dry ice for the time 0 data point. By 15, 30 and 60 minutes’ interval, tissue aliquots on room temperature have been put back on dry ice one by one. After all the tissue aliquots were placed back on dry ice, RNA extraction started immediately.

### RNA extraction and sequencing

Total RNA was extracted from each time point following molecular degradation off ice using

~100 mg of tissue using the RNeasy Lipid Tissue Mini kit (Qiagen) automatically by QIAcube according to the manufacturer’s protocol. RIN were measured by Agilent 2100 Bioanalyzer. For the RiboZero samples, sequencing libraries were constructed from total RNA using Illumina TruSeq Stranded Total RNA Ribo-Zero sample Prep Kit following the manufacturer’s protocol. For poly-A+ library, polyA-containing RNA molecules were purified from 1 µg DNAse treated total RNA and sequencing libraries were constructed using the Illumina TruSeq© RNA Sample Preparation v2 kit following the manufacturer’s protocol. The products were then purified and enriched with PCR to create the final cDNA library for high throughput sequencing using an Illumina HiSeq 2000 with paired end 2x100bp reads.

### RNA sequencing data processing

The Illumina Real Time Analysis (RTA) module performed image analysis, base calling, and the BCL Converter (CASAVA v1.8.2), generating FASTQ files containing the sequencing reads. These reads were aligned to the human genome (UCSC hg19 build) using the spliced-read mapper TopHat (v2.0.13) using the reference transcriptome to initially guide alignment, based on known transcripts in the Illumina iGenomes version of UCSC knownGene GTF file (using the “–G” argument in the software)^34^. Gene counts were generated using the featureCounts tool^35^ based on the more recent Ensembl v75, which was the last stable release for the hg19 genome build, using single end read counting [featureCounts –a $GTF –o $OUT $BAM]. There were 57,773 genes on the autosomes, sex chromosomes, and chrM. We converted counts to RPKM values using the total number of aligned reads across the autosomal and sex chromosomes (dropping reads mapping to the mitochondria chromosome).

### Public data processing

*Single cell RNAseq:* raw reads were downloaded and aligned directly from SRA using HISAT version 0.1.6-beta^36^. As above, featureCounts^35^ was used to quantify expression of genes relative to Ensembl v75.

*Blood:* Raw RNA-seq reads from all samples were downloaded from SRA and aligned to the genome using TopHat2^34^ (version 2.0.14) using the iGenomes transcriptome and genome annotations based on hg19. As above, featureCounts^35^ was used to quantify expression of genes relative to Ensembl v75.

*RNase*: Raw RNA-seq reads from all brain RNA in ABRF^22^ were downloaded from SRA and aligned to the genome using TopHat2^34^ (version 2.0.14) using the iGenomes transcriptome and genome annotations based on hg19. As above, featureCounts^35^ was used to quantify expression of genes relative to Ensembl v75.

*CommonMind Consortium (CMC)*: 547 BAM files were downloaded from Synapse, which were aligned with TopHat2 (version 2.0.9) using Ensembl v70 transcriptome annotation and the hg19 genome. As above, featureCounts^35^ was used to quantify expression of genes relative to Ensembl v75.

*GTEx*: Raw RNA-seq reads were downloaded from SRA and aligned to the genome using TopHat2^34^ (version 2.0.14) using the iGenomes transcriptome and genome annotations based on hg19. As above, featureCounts^35^ was used to quantify expression of genes relative to Ensembl v75

### Degradation data analysis

For the 20 samples in each library type, we separately modeled expression as a function of degradation time, adjusting for the donor, using the limma R Bioconductor package^37^.

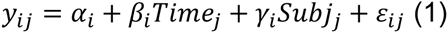

Expression values *y* for gene *i* and sample *j* were log_2_-transformed RPKMs values with an offset of 1, e.g. log2(RPKM+1). Resulting moderated T-statistics and p-values for the degradation time effect (e.g. *β*_*i*_) were obtained for each of the 57,773 genes for each library type. We also created degradation statistics for an overall degradation effect across all 40 samples by modeling expression as a function of time, adjusting for subject and library type, which we evaluated in microarray datasets below:

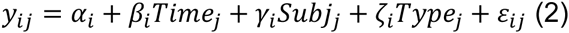

P-values were controlled for multiple testing using the Benjamini-Hochberg^38^ (B-H) procedure to control the false discovery rate (FDR), using the p-values from the expressed genes in each library type at mean RPKM > 0.05 (RiboZero: N=26,844, polyA: N=24,204, both: N=27,402). Gene set enrichment analyses were performed on the ordered degradation T-statistics from the polyA+ and RiboZero library types among those genes with Entrez Gene IDs using the `gseGO` and `gseKEGG` functions in the clusterProfiler R package^39^. Enrichment p-values were based on 10,000 permutations and subsequently adjusted for multiple testing using the B-H procedure to control the FDR.

### Cell type dataset analysis

We filtered the processed single cell RNA-seq data to the 28,363 expressed genes (at mean RPKM > 0.05) and 285 cells from adult donors that were previously classified as astrocytes, endothelial cells, microglia, neurons, oligodendrocytes, and oligodendrocyte progenitor cells (OPCs)^20^. We first modeled log_2_(RPKM+1) transformed expression as a function of cell type, and generated resulted F-statistics for the overall cell type effect at each gene. We then identified the most cell type-specific genes by identifying the set of genes that were more highly expressed in each cell type compared to all other cell types using t-tests. For each cell type versus all others, we selected those genes with p-values less than 10^−5^ (to ensure genes were differentially expressed) and then sorted by the log_2_ expression change and selected up to the top 250 cell type-specific genes. We then assigned each gene to one of the cell types (dropping 79 genes that were specific to two cell types) with the vast majority of genes not assigned to a cell type for each library type. We performed linear regression analysis to determine how degradation statistics differed among those genes for each cell type compared to non-cell type-specific genes, adjusting for mean expression of the gene.

### Blood dataset analysis

We performed analogous linear modeling for degradation time in the blood degradation RNA-seq dataset^21^, e.g. Eqn 1 above. To create a dataset with induced degradation confounding, we removed the 0 hour time point for individual 1 and the 84 hour time point for individual 2, and compared expression differences between the two individuals using linear modeling. The different statistical models included exprs ~ Individual, exprs ~ Individual + RIN, and exprs ~ Individual + Time, and we largely compared the resulting regression coefficients on Individual.

### LIBD discovery dataset modeling

We used the LIBD DLPFC polyA+ RNA-seq on 155 SCZD cases and 196 controls (criteria: ages between 17-80, gene assignment rate > 0.5, mapping rate > 0.7, RIN > 6, not outlying on 2nd ancestry PC, only self-reported Caucasians and African Americans) described in the companion Jaffe et al 2016. We fit several statistical models at the gene-level, modeling log_2_ transformed gene-level RPKM (with an offset of 1) as a function of:

1. Univariate: SCZD diagnosis
2. RIN-adjusted: SCZD diagnosis, adjusting for RIN
3. RIN2-adjusted: SCZD diagnosis, adjusting for RIN and RIN^2^
4. Full-adjusted: SCZD diagnosis, adjusting for age, sex, ancestry (SNP PCs 1, 5, 6, 9, 10, which were at least marginally associated with diagnosis), and then observed measures related to RNA quality: RIN, mitochondrial mapping rate, and gene assignment rate.
5. qSVA: SCZD diagnosis adjusting for “Full-adjusted” model (4) as well as the first 12 PCs from the degradation matrix (see below) based on DLPFC degradtion polyA+ libraries (selected using to using the BE algorithm^25^ in the sva Bioconductor package^26^ while providing the adjusted model as input).
6. PCA: SCZD diagnosis adjusting for “Adjusted” model as well as the first 23 PCs from the log_2_(RPKM+1) values (selected using to using the BE algorithm^25^ in the sva Bioconductor package^26^ while providing the adjusted model as input).
7. SVA: SCZD diagnosis adjusting for “Adjusted” model as well as 24 surrogate variables (SVs) from the log_2_(RPKM+1) values (selected using to using the BE algorithm^25^ in the sva Bioconductor package^26^ while providing the adjusted model as input).
8. Blood-qSVA: SCZD diagnosis adjusting for “Full-adjusted” model (4) as well as the first 12 PCs from the degradation matrix (see below) based on blood degradation polyA+ libraries (selected using to using the BE algorithm^25^ in the sva Bioconductor package^26^ while providing the adjusted model as input).

We used the `lmTest` and `ebayes` functions in the limma Bioconductor package^40^ to fit all of the statistical models to estimate log_2_ fold changes, moderated T-statistics, and corresponding p-values. Multiple testing correction via the false discovery rate (FDR) was applied using the set of 24,122 expressed genes (with mean RPKM > 0.1).

### qSVA approach

*Identifying degradation-susceptible regions:* We used the derfinder^24^ R Bioconductor package to identify expressed regions (ERs) for each library type (polyA+ and RiboZero) requiring a five read cutoff after normalizing each sample to 80M aligned reads. This resulted in 430,474 ERs across the 20 polyA+ samples and 2,435,196 ERs across the RiboZero samples. We first filtered the ERs for each library type to retain only those longer than 50 basepairs (bps), resulting in 214,304 ERs across the 20 polyA+ samples and 892,124 ERs across the RiboZero samples. The majority of ERs in the polyA+ samples were annotated to strictly exonic regions (58.9%) while the majority of ERs in the RiboZero samples were annotated to strictly intronic regions (69.5%). To identify the most degradation-susceptible regions in each library type, we fit the model in Eqn 1 above to the log_2_-adjusted read coverage (with an offset of 1) for each library type. As Bonferroni significance, there were 35,287 ERs and 515 ERs significantly associated with degradation time in the polyA+ and RiboZero datasets, respectively. We exported the top 1000 most significantly degradation-associated ERs (sorted by p-value) for the polyA+ data and the 515 significant ERs for the RiboZero data as BED files which represent the “degradation-susceptible regions”. These BED files are provided in Data S1 and S2, and also available at: https://github.com/nello/region_matrix. In the blood degradation dataset, we performed an analogous ER-level analysis – identifying ERs above 5 normalized reads, retaining those ERs > 50bp, and fitting Eqn 1 – to identify the 1000 most degradation-susceptible regions in blood (Data S3).

*Measuring the degradation matrix:* the “degradation matrix” is defined as the library-size normalized coverage across each of these regions in a user-provided dataset. There are several methods for calculating this degradation matrix.

- Directly from bigWig files: we used the `bwtool` tool to calculate the sum of read coverage^41^. Example code: `bwtool summary $BED $BW $OUT -header -fill=0 -with-sum` where $BED is either the polyA+ or RiboZero degradation-susceptible region BED file, $BW is the bigWig file for one sample, and $OUT is an output path.
- Directly from BAM files: We have provided a python script for calculating this degradation matrix directly from BAM files called “region matrix” at: https://github.com/nellore/region_matrix. Example code: `python region_matrix.py --regions sorted_polyA_degradation_regions_v2.bed -m bamAndSample_test.txt --wiggletools ~/bin/wiggletools `
- Directly from “Full coverage” derfinder^24^ objects using the `getRegionCoverage` function. Example code: ```
- gr = rtracklayer::import(“sorted_polyA_degradation_regions_v2.bed”) fullCov=fullCoverage(…) getRegionCoverage(fullCov, regions=gr, totalMapped = totalMapped) ```

These different implementations of the degradation matrix can be imported using the `sva` package and normalized to a common library size with the `read.degradation.matrix()` function.

*Correcting for RNA quality*: PCA is performed on the log_2_-transformed library size-normalized degradation matrix (with an offset of 1) and the top *K* PCs are selected, for example using the BE algorithm^25^. The set of these PCs are referred to as quality surrogate variables (qSVs), and can be included as adjustment variables in subsequent differential expression analyses. The estimation of the qSVs is implemented in the `qsva()` function in the `sva` package.

### CMC replications dataset analysis

We performed differential expression analysis on 159 patients and 172 controls (selecting on: total gene assignment rate > 0.3, alignment rate > 0.8, RIN > 6, ages between 18-80, non-outlying on genetic ancestry PCs 3 and 5 and keeping only reported Caucasians and African Americans). We similarly fit four of the statistical models at the gene-level, modeling log_2_ transformed gene-level RPKM (with an offset of 1) as a function of

1. Adjusted model: SCZD diagnosis, adjusting for age, sex, race, brain bank, RIN, gene assignment rate, alignment rate.
2. qSVA model: SCZD diagnosis adjusting for the “adjusted” model above and 14 qSVs based the degradation matrix constructed using the 515 regions based on the RiboZero.
3. PC adjustment: SCZD diagnosis adjusting for the “adjusted” model above and the first 27 PCs
4. SVA adjustment: PC adjustment: SCZD diagnosis adjusting for the “adjusted” model above and the 26 estimated SVs.

In these replication data we did not perform FDR correction. We were using the study for replication, not discovery, and therefore only used the features that were expressed in the LIBD discovery dataset regardless of the expression levels in CMC. We considered features replicated if they had the same directionality for the SCZD versus control log_2_ fold change and were marginally significant (at p < 0.05). Gene body coverage bias were measured using the `geneBody_coverage.py` script in the RSeQC package^42^ and the provided housekeeping genes. The coverage bias was summarized by the first PC of the percentiles of read coverage across the transcripts, which captured >75% of variability in each dataset.

### GTEx analysis

We retained all GTEx samples that had RINs > 5 and belonged to sub-tissues (`SMTSD` metadata column) with at least 40 samples, resulting in data on 9,502 samples across 49 sub-tissues. We retained the 36,552 genes that had mean RPKM > 0.2 in at least one sub-tissue. We modeled differential expression of each of 48 sub-tissues compared to “Brain - Frontal Cortex (BA9)”, and measured the Pearson correlation present in the resulting DEqual plots, e.g. between these sub-tissue specific log_2_ fold changes to the DLPFC polyA+ degradation data log_2_ fold changes for degradation time (from Eqn 1 above). We compared the DEqual correlations to the difference in chrM mapping rates using the `cor.test()` function in R within all, and also just brain-specific, sub-tissues.

### Microarray data processing and analysis of published studies

We sought to leverage the degradation DLPFC RNA-seq data to correct previously published microarray studies for brain disorders, as there are few large publicly-available RNA-seq datasets for disorders other than schizophrenia. We extrapolated the expression levels of the probes for each microarray platform in our degradation RNA-seq dataset using the following approach:

A. Map probe sequences to the transcriptome + genome using TopHat2 and the iGenomes annotation files
B. Filter resulting alignments to remove those probes that were multi-mapped, contained indels, mapped to the sex chromosomes and chrM. Resulting alignments with splice junctions, e.g. those probes that crossed exon-exon splice junctions were retained.
C. Extract coverage of the probe alignments in the degradation RNA-seq data using the `getRegionCoverage` function in the derfinder package, including those spliced alignments
D. Fit Eqn 2 above in extrapolated coverage data from the degradation RNA-seq data to identify the likely probe susceptibility to degradation.

We considered the Illumina HumanRef-8 v3.0 (GPL6883) platform from Voineagu, et al.^27^ and the Rosetta/Merck Human 44k 1.1 (GPL4372) platform from Zhang, et al.^28^. There were 22,466 and 33,833 probes that resulted from steps A-D above for GPL6883 and GPL4372 respectively, of which 9,340 and 7,716 were associated with degradation time when extrapolated to the RNA-seq degradation data at FDR < 5%, respectively. We selected the more stringent set of 1,129 and 653 probes most associated with degradation at Bonferroni < 1% significance for the GPL6883 and GPL4372 platforms, respectively to assess potential enrichment for degradation confounding. For the ASD dataset, we considered univariate statistical models for diagnosis (there were no probes differentially expressed at FDR < 5%), and for the AD dataset, we considered covariate-adjusted (there were 17,565 probes at FDR < 5%)

## Acknowledgements

The Genotype-Tissue Expression (GTEx) Project was supported by the Common Fund of the Office of the Director of the National Institutes of Health. Additional funds were provided by the NCI, NHGRI, NHLBI, NIDA, NIMH, and NINDS. Donors were enrolled at Biospecimen Source Sites funded by NCI\SAIC-Frederick, Inc. (SAIC-F) subcontracts to the National Disease Research Interchange (10XS170), Roswell Park Cancer Institute (10XS171), and Science Care, Inc. (X10S172). The Laboratory, Data Analysis, and Coordinating Center (LDACC) was funded through a contract (HHSN268201000029C) to The Broad Institute, Inc. Biorepository operations were funded through an SAIC-F subcontract to Van Andel Institute (10ST1035). Additional data repository and project management were provided by SAIC-F (HHSN261200800001E). The Brain Bank was supported by a supplement to University of Miami grants DA006227 & DA033684 and to contract N01MH000028. Statistical Methods development grants were made to the University of Geneva (MH090941 & MH101814), the University of Chicago (MH090951, MH090937, MH101820, MH101825), the University of North Carolina - Chapel Hill (MH090936 & MH101819), Harvard University (MH090948), Stanford University (MH101782), Washington University St Louis (MH101810), and the University of Pennsylvania (MH101822). The data used for the analyses described in this manuscript were obtained from dbGaP accession number phs000424.v6.p1 on October 6, 2015.

Data were generated as part of the CommonMind Consortium supported by funding from Takeda Pharmaceuticals Company Limited, F. Hoffman-La Roche Ltd and NIH grants R01MH085542, R01MH093725, P50MH066392, P50MH080405, R01MH097276, RO1-MH-075916, P50M096891, P50MH084053S1, R37MH057881 and R37MH057881S1, HHSN271201300031C, AG02219, AG05138 and MH06692. Brain tissue for the study was obtained from the following brain bank collections: the Mount Sinai NIH Brain and Tissue Repository, the University of Pennsylvania Alzheimer’s Disease Core Center, the University of Pittsburgh NeuroBioBank and Brain and Tissue Repositories and the NIMH Human Brain Collection Core. CMC Leadership: Pamela Sklar, Joseph Buxbaum (Icahn School of Medicine at Mount Sinai), Bernie Devlin, David Lewis (University of Pittsburgh), Raquel Gur, Chang-Gyu Hahn (University of Pennsylvania), Keisuke Hirai, Hiroyoshi Toyoshiba (Takeda Pharmaceuticals Company Limited), Enrico Domenici, Laurent Essioux (F. Hoffman-La Roche Ltd), Lara Mangravite, Mette Peters (Sage Bionetworks), Thomas Lehner, Barbara Lipska (NIMH).

**Supplementary Information:** supplementary table, figure, and data legends are provided at the end of the paper.

## Author Contributions

A.E.J – conceived the study, designed and oversaw the research project, performed primary data processing and analyses, led the writing of the manuscript

R.T. – performed tissue degradation experiment and extracted RNA

A.L.N – performed cell-type specificity analyses

M.K. – performed microarray-to-RNAseq bioinformatics analysis

A.N. – wrote software for computing coverage of degradation-susceptible regions

Y.J. – oversaw RNA sequencing data generation (library preparation, and sequencing)

T.M.H., J.E.K. – provided brain tissue for degradation experiment, contributed to writing of the manuscript.

R.E.S. – contributed to experimental design

J.T.L. – contributed to qSVA algorithm development

D.R.W. – contributed to interpretation of the analyses and writing of the manuscript

**Author information:** sequencing reads are available through SRA at accession numbers: [**TBD].** Reprints and permissions information is available at https://www.nature.com/reprints. Correspondence and requests for materials should be addressed to Andrew Jaffe. The authors have no competing interests to report.

## Supplementary Figure Legends

Figure S1: Principal component analysis (PCA) on gene expression levels [log_2_(RPKM+1) scale] suggests that degradation time is the largest component of variability in both (A) polyA+ and (B) RiboZero RNA-seq sequencing libraries. Each line represents samples from the same subject. The p-values are based on linear regression of each top PC on degradation time, adjusting for the subject.

Figure S2: Comparisons of RNA degradation with cell type-specificity of genes by library type. The genes more influenced by cell type from Darmanis et al^20^ single cell RNA-seq data (via an F-statistic for an ANOVA of the cell type effect) were more likely to be decreased by degradation time in the frontal cortex in the (A) RiboZero and (B) polyA+ library types.

Figure S3: Comparisons between degradation effects (via T-statistics) comparing brain tissue, blood, and brain-derived RNA. Globally, there was (A) little overlap between blood-based and brain tissue-based degradation effects on gene expression levels and (B) high overlap between leaving brain tissue at room temperature compared to RNase A-treated brain-derived RNA. Each point is a gene.

Figure S4: Analogous DEqual plots using log_2_ fold changes for the schizophrenia and degradation effects rather than the T-statistics shown in Figure 1. (A) DEqual plot for univariate case-control analysis shows strong correlation between degradation and diagnosis effects, (B) DEqual plot for RIN-adjusted case-control differences largely fails to remove degradation bias, and (C) DEqual plot when adjusting for observed clinical and technical covariates, including age, sex, ethnicity, chrM mapping rate, gene assignment rate, and RIN, also fail to remove degradation bias.

Figure S5: Modeling RIN to further allowing for non-linear effects (RIN + RIN^2^) shows little difference in (A) removing positive correlation from the T-statistic DEqual plot and (B) comparing T-statistics for diagnosis to the RIN-adjusted model (y-axis).

Figure S6: Associating estimated quality surrogate variables (qSVs) with observed covariates. Color associated with the –log10 p-value from regression modeling of each qSV versus each covariate in the LIBD discovery dataset in polyA+ data. “geneBodyBiasPc”: the first PC of the percentiles of read coverage from 3’ to 5’; “snpPC1”: the first principal component of genomic ancestry via genetic SNP data, “totalAssignedGene”: the proportion of aligned reads that were assigned to genes during read counting; “mitoRate”: the proportion of reads that aligned to chrM; “PMI”: post-mortem interval, “Dx”: diagnosis, SCZD or control.

Figure S7: Associating estimated quality surrogate variables (qSVs) with observed covariates. Color associated with the –log10 p-value from regression modeling of each qSV versus each covariate in the CMC replication dataset in RiboZero data. “geneBodyBiasPc”: the first PC of the percentiles of read coverage from 3’ to 5’; “snpPC1”: the first principal component of genomic ancestry via genetic SNP data, “totalAssignedGene”: the proportion of aligned reads that were assigned to genes during read counting; “mitoRate”: the proportion of reads that aligned to chrM; “PMI”: post-mortem interval, “Dx”: diagnosis, SCZD or control. Other details of covariates are provided from Synapse.org

Figure S8: T-statistics DEqual plot demonstrating that principal component analysis (PCA) on the entire transcriptome also successfully removes positive correlation between degradation and SCZD effects.

Figure S9: T-statistics DEqual plot demonstrating that surrogate variable analysis (SVA) on the entire transcriptome also fails to removes positive correlation between degradation and SCZD effects.

Figure S10: The DLPFC-derived degradation-susceptible regions associate with degradation effects in the (A) blood and (B) ABRF RNase A degradation datasets. These PCs would be considered the first quality surrogate variables (qSVs) in the qSVA framework.

Figure S11: The qSVA approach reduces degradation bias in blood. (A) DEqual plot for the qSV-adjusted individual effects versus degradation effects shows no correlation. Statistical bias for the (B) qSVA versus the (C) RIN adjustment approaches in the degradation-confounding design in the blood degradation dataset. Y-axis: “bias” is the difference in log_2_ fold changes comparing each model to the “true” adjustment model, which is adjusting for degradation time of each sample. X-axis: the average log_2_ fold change for the time- and either qSVA- or RIN-adjusted models. Each point is a gene.

Figure S12: Comparing brain-derived qSVs from blood-derived qSVs suggests weak residual RNA degradation biases. (A) The DEqual plot (on log_2_ fold changes) shows stronger negative correlation between diagnosis and degradation effects. (B) Moderately correlated log_2_ fold change estimates for the SCZD effects across the two qSVA implementations.

Figure S13: Comparing the ability of the qSVA framework to remove degradation bias in non-brain tissues using brain-derived degradation regions. (A) Strong correlation between the correlation in DEqual plots for univariate tissue-specific expression effects and DLPFC degradation compared to the difference in chrM mapping rates (which was the variable most associated with degradation time in brain data, see Table S2). This significant correlation was driven by the brain (blue) samples as the correlation was not significant in the non-brain samples (red). (B) No correlation between the DEqual summary statistic and the difference in chrM mapping rates in either brain or non-brain tissues following qSVA correction using DLPFC-derived degradation regions.

## Supplementary Table Legends

Table S1: RIN values for DLPFC degradation experiment

Table S2: Associations between degradation time and sample- and library-specific covariates following linear regression analysis of degradation time on each covariate, adjusting for donor.

Table S3: Degradation susceptibility statistics at each gene in Ensembl v75. These statistics are the y-axis of the DEqual plots. Prefixes: “polyA” – the 20 polyA+ samples, “Ribo” – the 20 RiboZero samples, “both” – the combined 40 samples. Suffixes: “timeFC” – the log2 fold change in expression per minute of degradation from regression modeling, adjusting for donor, “Tstat” – moderated T-statistic from limma for the standardized “timeFC”, “Pvalue” – corresponding p-value for moderated T-statistic. “meanExprs” – the mean expression levels across the different samples in the RPKM scale. “FDR” were based on the B-H correction procedure on those genes with RPKM > 0.1 in each sample set (NA means expression was below the threshold).

Table S4: Gene set enrichment results for GO and KEGG for those sets with FDR < 5% in both polyA+ and RiboZero library types in the DLPFC degradation datasets, based on the T-statistics for degradation effects.

Table S5: Comparison of cell type specificity of degradation effects by library and cell type.

## References

1 Cancer Genome Atlas Research, N. Integrated genomic analyses of ovarian carcinoma. Nature 474, 609–615, doi:10.1038/nature10166 (2011).

2 Lappalainen, T. et al. Transcriptome and genome sequencing uncovers functional variation in humans. Nature 501, 506–511, doi:10.1038/nature12531 (2013).

3 Sirota, M. et al. Discovery and preclinical validation of drug indications using compendia of public gene expression data. Science translational medicine 3, 96ra77, doi:10.1126/scitranslmed.3001318 (2011).

4 Colantuoni, C. et al. Temporal dynamics and genetic control of transcription in the human prefrontal cortex. Nature 478, 519–523, doi:10.1038/nature10524 (2011).

5 Kang, H. J. et al. Spatio-temporal transcriptome of the human brain. Nature 478, 483–489, doi:10.1038/nature10523 (2011).

6 Edgar, R., Domrachev, M. & Lash, A. E. Gene Expression Omnibus: NCBI gene expression and hybridization array data repository. Nucleic acids research 30, 207–210 (2002).

7 Irizarry, R. A. et al. Exploration, normalization, and summaries of high density oligonucleotide array probe level data. Biostatistics 4, 249–264, doi:10.1093/biostatistics/4.2.249 (2003).

8 Du, P., Kibbe, W. A. & Lin, S. M. lumi: a pipeline for processing Illumina microarray. Bioinformatics 24, 1547–1548, doi:10.1093/bioinformatics/btn224 (2008).

9 Hansen, K. D., Irizarry, R. A. & Wu, Z. Removing technical variability in RNA-seq data using conditional quantile normalization. Biostatistics 13, 204–216, doi:10.1093/biostatistics/kxr054 (2012).

10 Risso, D., Ngai, J., Speed, T. P. & Dudoit, S. Normalization of RNA-seq data using factor analysis of control genes or samples. Nature biotechnology 32, 896–902, doi:10.1038/nbt.2931 (2014).

11 Leek, J. T. et al. Tackling the widespread and critical impact of batch effects in high-throughput data. Nature reviews. Genetics 11, 733–739, doi:10.1038/nrg2825 (2010).

12 Li, S. et al. Detecting and correcting systematic variation in large-scale RNA sequencing data. Nature biotechnology 32, 888–895, doi:10.1038/nbt.3000 (2014).

13 Wang, C. et al. The concordance between RNA-seq and microarray data depends on chemical treatment and transcript abundance. Nature biotechnology 32, 926–932, doi:10.1038/nbt.3001 (2014).

14 Consortium, S. M.-I. A comprehensive assessment of RNA-seq accuracy, reproducibility and information content by the Sequencing Quality Control Consortium. Nature biotechnology 32, 903–914, doi:10.1038/nbt.2957 (2014).

15 t Hoen, P. A. et al. Reproducibility of high-throughput mRNA and small RNA sequencing across laboratories. Nature biotechnology 31, 1015–1022, doi:10.1038/nbt.2702 (2013).

16 Adiconis, X. et al. Comparative analysis of RNA sequencing methods for degraded or low-input samples. Nature methods 10, 623–629, doi:10.1038/nmeth.2483 (2013).

17 Schroeder, A. et al. The RIN: an RNA integrity number for assigning integrity values to RNA measurements. BMC molecular biology 7, 3, doi:10.1186/1471-2199-7-3 (2006).

18 Consortium E. R. REMC Standards and Guidelines for RNA-sequencing. (2014).

19 Wang, L. et al. Measure transcript integrity using RNA-seq data. BMC bioinformatics 17, 58, doi:10.1186/s12859-016-0922-z (2016).

20 Darmanis, S. et al. A survey of human brain transcriptome diversity at the single cell level. Proceedings of the National Academy of Sciences of the United States of America 112, 7285–7290, doi:10.1073/pnas.1507125112 (2015).

21 Gallego Romero, I., Pai, A. A., Tung, J. & Gilad, Y. RNA-seq: impact of RNA degradation on transcript quantification. BMC biology 12, 42, doi:10.1186/1741-7007-12-42 (2014).

22 Li, S. et al. Multi-platform assessment of transcriptome profiling using RNA-seq in the ABRF next-generation sequencing study. Nature biotechnology 32, 915–925, doi:10.1038/nbt.2972 (2014).

23 Leek, J. T. & Storey, J. D. Capturing heterogeneity in gene expression studies by surrogate variable analysis. PLoS genetics 3, 1724–1735, doi:10.1371/journal.pgen.0030161 (2007).

24 Collado Torres, L. et al. Flexible expressed region analysis for RNA-seq with derfinder. bioRxiv, doi:10.1101/015370 (2016).

25 Buja, A. & Eyuboglu, N. Remarks on Parallel Analysis. Multivariate Behavioral Research 27, 509–540, doi:10.1207/s15327906mbr2704_2 (1992).

26 Leek, J. T., Johnson, W. E., Parker, H. S., Jaffe, A. E. & Storey, J. D. The sva package for removing batch effects and other unwanted variation in high-throughput experiments. Bioinformatics 28, 882–883, doi:10.1093/bioinformatics/bts034 (2012).

27 Voineagu, I. et al. Transcriptomic analysis of autistic brain reveals convergent molecular pathology. Nature 474, 380–384, doi:10.1038/nature10110 (2011).

28 Zhang, B. et al. Integrated systems approach identifies genetic nodes and networks in late-onset Alzheimer's disease. Cell 153, 707–720, doi:10.1016/j.cell.2013.03.030 (2013).

29 Jaffe, A. E. Postmortem human brain genomics in neuropsychiatric disorders--how far can we go? Current opinion in neurobiology 36, 107–111, doi:10.1016/j.conb.2015.11.002 (2016).

30 Jaffe, A. E. et al. Mapping DNA methylation across development, genotype and schizophrenia in the human frontal cortex. Nat Neurosci 19, 40–47, doi:10.1038/nn.4181 (2016).

31 Jaffe, A. E. et al. Developmental regulation of human cortex transcription and its clinical relevance at single base resolution. Nat Neurosci 18, 154–161, doi:10.1038/nn.3898 (2015).

32 Jaffe, A. E. & Irizarry, R. A. Accounting for cellular heterogeneity is critical in epigenomewide association studies. Genome Biol 15, R31, doi:10.1186/gb-2014-15-2-r31 (2014).

## References

33 Lipska, B. K. et al. Critical factors in gene expression in postmortem human brain: Focus on studies in schizophrenia. Biol Psychiatry 60, 650–658, doi:10.1016/j.biopsych.2006.06.019 (2006).

34 Kim, D. et al. TopHat2: accurate alignment of transcriptomes in the presence of insertions, deletions and gene fusions. Genome Biol 14, R36, doi:10.1186/gb-2013-14-4-r36 (2013).

35 Liao, Y., Smyth, G. K. & Shi, W. featureCounts: an efficient general purpose program for assigning sequence reads to genomic features. Bioinformatics 30, 923–930, doi:10.1093/bioinformatics/btt656 (2014).

36 Kim, D., Langmead, B. & Salzberg, S. L. HISAT: a fast spliced aligner with low memory requirements. Nature methods 12, 357–360, doi:10.1038/nmeth.3317 (2015).

37 Smyth, G. K. Linear models and empirical bayes methods for assessing differential expression in microarray experiments. Statistical applications in genetics and molecular biology 3, Article3, doi:10.2202/1544-6115.1027 (2004).

38 Benjamini, Y. & Hochberg, Y. Controlling the False Discovery Rate: A Practical and Powerful Approach to Multiple Testing. Journal of the Royal Statistical Society. Series B (Methodological) 57, 289–300, doi:10.2307/2346101 (1995).

39 Yu, G., Wang, L. G., Han, Y. & He, Q. Y. clusterProfiler: an R package for comparing biological themes among gene clusters. Omics: a journal of integrative biology 16, 284–287, doi:10.1089/omi.2011.0118 (2012).

40 Smyth, G. K. Linear models and empirical Bayes methods for assessing differential expression in microarray experiments. Statistical applications in genetics and molecular biology 3, Article 3 (2004).

41 Pohl, A. & Beato, M. bwtool: a tool for bigWig files. Bioinformatics 30, 1618–1619, doi:10.1093/bioinformatics/btu056 (2014).

42 Wang, L., Wang, S. & Li, W. RSeQC: quality control of RNA-seq experiments. Bioinformatics 28, 2184–2185, doi:10.1093/bioinformatics/bts356 (2012).

